# Using spatiotemporal source separation to identify prominent features in multichannel data without sinusoidal filters

**DOI:** 10.1101/190470

**Authors:** Michael X Cohen

## Abstract

The number of simultaneously recorded electrodes in neuroscience is steadily increasing, providing new opportunities for understanding brain function, but also new challenges for appropriately dealing with the increase in dimensionality. Multivariate source-separation analysis methods have been particularly effective at improving signal-to-noise ratio while reducing the dimensionality of the data, and are widely used for cleaning, classifying, and source-localizing multichannel neural time series data. Most source-separation methods produce a spatial component (that is, a weighted combination of channels to produce one time series); here, this is extended to apply source-separation to a time series, with the idea of obtaining a weighted combination of successive time points, such that the weights are optimized to satisfy some criteria. This is achieved via a two-stage source-separation procedure, in which an optimal spatial filter is first constructed, and then its optimal temporal basis function is computed. This second stage is achieved with a time-delay-embedding matrix, in which additional rows of a matrix are created from time-delayed versions of existing rows. The optimal spatial and temporal weights can be obtained by solving a generalized eigendecomposition of covariance matrices. The method is demonstrated in simulated data and in an empirical EEG study on theta-band activity during response conflict. Spatiotemporal source separation has several advantages, including defining empirical filters without the need to apply sinusoidal narrowband filters.

---

Population neural activity measured through the electroencephalogram (EEG) or local field potential (LFP) is often rhythmic (Buzsáki and Draguhn, 2004). These rhythms can be grouped into bands according to dominant frequency characteristics (e.g., delta, theta, alpha, gamma), and reflect oscillations in population-level excitability (Wang, 2010). Neural oscillations have been linked to myriad neural and cognitive functions over the past century. Although questions remain regarding the precise origins and computational roles of oscillations, it is undeniable that neural oscillations are robust markers of neurocognitive phenomena and can be used to link findings across species and spatial scales (Klimesch, 1999; Le Van Quyen and Van Quyen, 2011; Buzsáki et al., 2013). Neural oscillations are also increasingly being linked to brain disorders ranging from Schizophrenia to Parkinson’s to depression to anxiety (Uhlhaas and Singer, 2010; Ba§ar, 2013; Oswal et al., 2013).

In some cases, neural oscillations can be identified solely by qualitative visual inspection (Cole and Voytek, 2017). However, these cases tend to be the exception rather than the rule. Instead, most investigations require signal processing methods before interpretations can be made. Signal processing is necessary because electrodes measure activity from multiple sources simultaneously, because the signal-to-noise characteristics of the data can be low, and because neural oscillations have nonlinearities such as bursting that can limit their detectability when using signal-processing methods that are optimized for stationary signals (e.g., the Fourier transform). Therefore, it is important to be able to identify neural oscillations in potentially noisy data with potentially weak signals.

The most commonly used analysis method for identifying neural oscillations is to apply temporal filters such as Morlet wavelets or narrowband FIR filters, and then extract estimates of time-varying power and phase values (Cohen, 2014a). However, such narrowband filters impose sinusoidality on time series data, thus biasing the results to identifying sinusoid-looking features of the data. It is therefore of theoretical and practical interest to be able to identify important temporal features of data without imposing any specific waveform shape on the results.

A promising approach for decomposing multichannel EEG data into different potential contributing sources is source-separation analyses, which have the goal of finding weighted combinations of activity across different electrodes, where the weights are defined according to some criteria, and where the linear weighted sum of activity across all electrodes is used to generate a single time series vector (the component time series). Depending on the goal of the analyses, the criteria can be anatomical (e.g., dipole fitting or distributed localization in minimum-norm or beamforming; (Hillebrand and Barnes, 2005), signal-distribution independence (e.g., independent components analysis; Jung et al., 2001), or contrasts between two features of the data (e.g., generalized eigendecomposition; Parra et al., 2005). Regardless, the result of the source separation is a component time series that comprises a linear combination of data at all electrodes. From here, researchers often apply standard temporal filters such as wavelets or narrowband FIR filters. This means that only the spatial features are optimized for different sources of variance, not the temporal features.

The purpose of this paper is to extend existing methods for identifying spatiotemporal features of multichannel electrophysiology data. The method involves combining source separation techniques with time-delay-embedding to identify prominent features of neural signals without the need to impose a sinusoidal filter. Instead, the optimal filter kernel is computed directly from the data (the filter is “optimal” in that it maximizes researcher-specified criteria). This multivariate approach facilitates separating multiple spatiotemporal sources, providing those sources have differentiable projections onto the recording electrodes. It is perhaps best suited for task-related data, in which one compares an experiment condition against a baseline time period or a baseline condition. The method is applied to simulated data and to empirical human EEG data. Issues of practical implementation are also discussed.

## Methods

### Broad overview of the spatiotemporal filter

The source separation method presented here involves two stages (see Figure 1 for a visual overview). First, an optimal spatial filter is constructed with the goal of reducing the dimensionality of the data from M channels to C components, where C<<M. Each component is a linear combination of all electrodes that maximizes some user-specified objective function (e.g., a comparison between conditions). The spatially filtered time series data are then time-delay-embedded, and a temporal source separation is applied to that delay-embedded matrix. The result of this second source separation is an empirically derived filter that can be applied to the data, to which time series analyses can be applied such as time-domain averaging or applying the Hilbert transform to extract power and phase estimates. Note that at no point in this procedure are narrowband temporal filters such as wavelets, FIR filters, or FFT-based filters imposed on the time series data.

**Figure 1.**
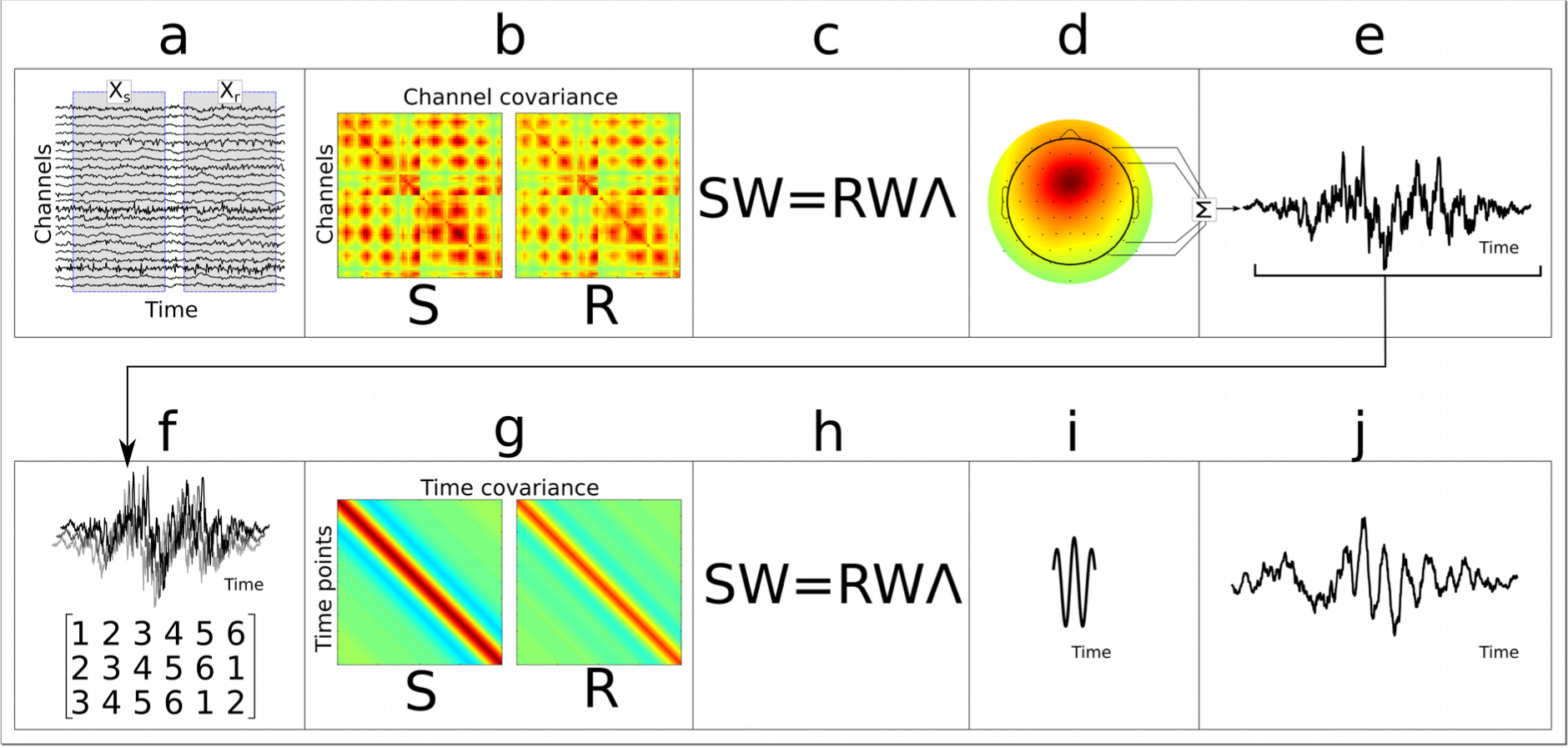
The ten-step procedure for obtaining the spatiotemporal filter. (a) Raw data showing two regions containing the signal of interest (**X**_s_) and reference (**X**_**r**_). (b) Channel covariance matrices are computed for each of these time windows, (c) which are then used in a generalized eigendecomposition. (d) A spatial filter is selected (column of **W**) and the filter forward model can be visualized as a topographical map. (e) The weighted combination of all electrodes is a time series (Hann-tapered here for visibility). Most source-separation methods stop at this step, but the important temporal features of this component time series can be better extracted via a second source separation stage. (f) The time series data are delay-embedded, which means new rows of the data matrix are created from delayed versions of the original row(s). (g) The time covariance matrices from time windows to be maximized (S) vs. minimized (**R**) are used to form two covariance matrices, (h) on which a generalized eigendecomposition is performed. The eigenvector with the largest eigenvalue (i) is the optimal basis vector that separates **S** from **R**, and is used as an empirically defined temporal filter kernel that can be applied to the data from panel e, which (j) creates the spatiotemporally filtered data. Note that steps a-e and steps f-j are the same except for the application to the spatial or temporal domains. Data in panels d, i, and j can be pooled and compared across individuals.

### Geometric and analytic explanations of the spatiotemporal filter

EEG data are often conceptualized as a mixture of electrical fields produced by several neural sources. Key to multivariate decomposition methods is the assumption that this mixture is linear because the electrical fields propagate simultaneously (within measurement capabilities) from all sources to all electrodes (Nunez and Srinivasan, 2006). Thus, the electrode-level data can be conceptualized as

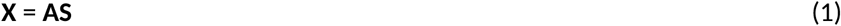

where **X** is the observed channels × time matrix, **S** is the underlying sources of activity, and **A** is a transformation matrix. **A** and **S** are *a priori* unknown, which presents a challenge for scientists.

Therefore, the goal of multivariate source-separation methods is to make assumptions about **A** and **S** in attempt to best estimate an appropriate **A**^-1^ that could left-multiply **X** to gain insights into **S**. These assumptions can be based on independence and non-Gaussian distributions (e.g., independent components analysis), or on frequency characteristics, anatomical locations, or differences between experiment conditions or time periods. The latter approach is taken here.

Geometrically, the recorded data in **X** occupy an M-dimensional space, where M is the number of electrodes. Each basis vector in this space is defined by each electrode, and each time point of data can be thought of as a point in this space, with the projection along each basis vector **i** equal to the microvolt value recorded at electrode M_i_.

The purpose of the first source separation stage is to find a better set of spatial basis vectors with the goal of reducing the dimensionality of the data from M to C, where C<<M (for convenience, C is often 1, but multiple sources can be extracted by allowing C>1, as will be shown in the second simulation below) and C is defined to optimize the multivariate power ratio between two experiment conditions, or between two time periods. Let the covariance matrices for the two conditions be matrix **S** (the “signal” to be maximized) and matrix **R** (the “reference” data):

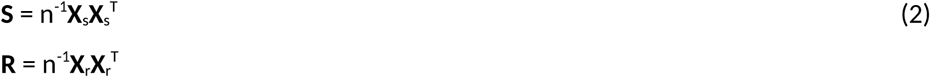

where **X** is the mean-centered channels × time (MxN) data matrix, ^T^ is the matrix transpose, n is the number of time points, and subscripts _s_ and _r_ indicate subsets of the data corresponding to time periods to maximize and minimize (signal vs. reference). For a trial-based experiment, the covariance matrices should be computed per trial and then averaged over all trials.

Finding a set of M weights (in vector **w**) such that the weighted sum of activity at all electrodes maximizes the distance (or power ratio) between **S** and **R** can be obtained by the Rayleigh Quotient:

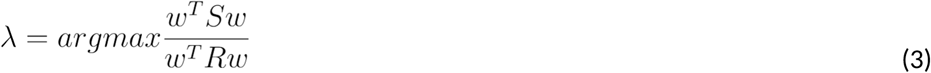

Note that w^T^**S**w is a single number representing the “energy” in matrix **S** along direction **w**. Therefore, the goal is to find the M×1 vector **w** such that along direction **w**, the ratio between **S** and **R** is maximized (*λ* is the value of that ratio). For a complete set of weights (M M × 1 vectors), this can be solved as

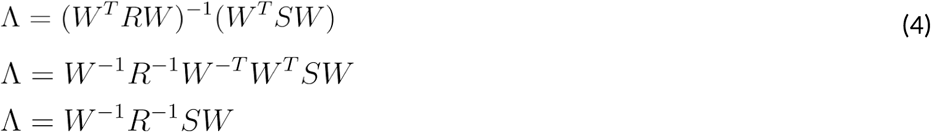

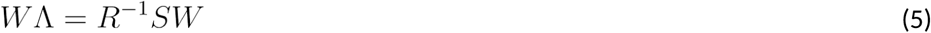

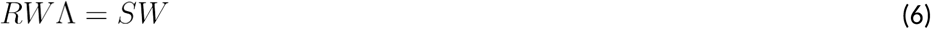

In other words, the problem of separating data represented by covariance matrices **S** and **R** can be solved via generalized eigenvalue decomposition. This insight has inspired many source separation applications in neuroscience (Parra et al., 2005; Tomé, 2006; Blankertz et al., 2008), and in this sense, the method presented here follows directly from this tradition. Equation 5 is perhaps a more intuitive way to conceptualize this mechanism of source separation (eigendecomposition of the matrix “division” **S**/**R**), although equation 6 is closer to the implementation in Matlab using `[W,L]=eig(S,R)` where M×M matrix **W** contains the eigenvectors in the columns, and M×M matrix **L** (this is ∧ in equation 6) contains the corresponding eigenvalues in the diagonal. The eigenvector associated with the largest eigenvalue is the basis vector that maximally separates **S** from **R**.

Note that although both **S** and **R** are symmetric positive semidefinite matrices, the eigendecomposition is implemented on the matrix product **R^-1^S**, which is not symmetric. Therefore, the eigenvectors in matrix **W** are not constrained to be orthogonal as they are in principal components analysis. In practical applications, data matrices are often positive semi-definite (not positive definite) because standard EEG preprocessing reduces the rank of the data. Reduced-rank matrices are not problematic for the method presented here because generally one is interested in only those components with the largest eigenvalues; eigenvectors with repeated or zero-valued eigenvalues can be ignored. The weights themselves mix suppressing irrelevant electrodes (or time points for the temporal separation stage described below) and boosting relevant electrodes, and therefore they can be difficult to interpret directly. Instead, the “forward model” (sometimes also called “activation pattern”) of the filter is interpreted and averaged across subjects, and is computed as (**SW**)(**W^T^SW**)^-1^ (this can be shortened to **W^-T^** for full-rank matrices) (Haufe et al., 2014). The important part of this formula is **Sw**, in other words, multiplying the covariance matrix by the eigenvector used to filter the data; the multiplication by (**W^T^SW**)^-1^ is a scaling factor.

Regularization was added as 0.1% of the variance to the diagonal of the **R** matrix as follows:

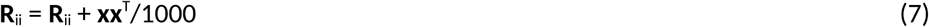

where **x** is the mean-centered time series data from channel **i** (as a row vector) and ^T^ is the vector transpose. Various levels of regularization were examined; this amount of regularization either improved slightly or did not appreciably affect the results. There are also several algorithms for regularization, including Thikonov, eigenvalue shrinkage, and so on. These were not systematically explored here, although it is likely that the small amount of regularization would not be appreciably different for different regularization methods.

Geometrically, one can think of the eigenvectors in **W** as providing a new set of basis vectors in the data space such that the basis vector defined by the column in **W** with the largest associated eigenvalue maximizes the power ratio between **S** and **R**. Projecting the channel data **X** onto the largest eigenvector (in practice, this is achieved by computing the weighted sum of all electrodes, or **y**=**w^T^X**) is the component time series that maximizes the researcher-specified criteria that were defined when creating matrices **S** and **R**.

Now the data have been reduced from dimensionality M to dimensionality C. That completes the first stage of the method. The second stage is to use that component to create a new multivariate space by time-delay-embedding the data, thus expanding the dimensionality to CD dimensions (where D is the number of delay embeds). In practice, it may be easier to delay-embed separately each component c∈C, thus creating C D-dimensional delay embedded matrices. Time-delay embedding means adding rows to a matrix that are defined by time-delayed versions of the original data (see Figure 1f).

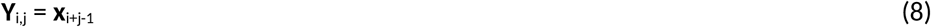

where **x** is the component time series vector (Figure 1e) and *i* and *j* correspond to row and column indices. This can be implemented using a for-loop or, because the delay-embedded matrix is a form of a Hankel matrix, the Matlab command `hankel`. Because subsequent time points are not redundant (assuming the number of embeds is less than the number of time points, which is generally the case for EEG data), matrix **Y** has a rank equal to its embedding dimension. The purpose of creating the delay-embedded matrix is to apply a source-separation decomposition on the time series data. The weights created for each row in the matrix reflect weights for successive time points. It thus follows that taking the weighted combination of the delayed time series is equivalent to applying a temporal filter to time series data. The main difference is that the temporal weights can be defined according to eigenvectors computed from the data, rather than, e.g., a Morlet wavelet that would be applied to the data for narrowband filtering. For example, a single embedding would produce a 2xN matrix, and row weights of [-1 1] would correspond to the first derivative of the time series. The number of embeds should be at least as large as the expected empirical filter kernel.

The geometric interpretation of this step is an expansion of the one-dimensional subspace identified in the first source-separation phase to a D-dimensional space in which each basis vector is defined by each time point. Thus, the purpose of this second source separation is to identify a new set of basis vectors in this space that maximizes the same researcher-specified criteria as described for the first stage. The primary difference is that instead of obtaining a spatial filter, these eigenvectors produce a temporal filter (based on the output of the optimally spatially filtered data).

The weighted combination of the delay-embedded data in matrix **Y** is a time series to which time series analyses can be applied. The primary analysis applied here is time-frequency analysis, implemented by taking the magnitude of the Hilbert transform of the time series.

The sign of an eigenvector is often not meaningful—the eigenvector points along a dimension; that dimension can be equally well indicated regardless of whether the vector points “forwards” or “backwards.” For visual clarity, the sign of the topographical maps was adjusted so that the electrode with the largest magnitude was forced to be positive (this is a common procedure in principal components analysis).

In theory, these two source separation stages could be implemented in one shot by delay-embedding the M-channel time series. However, this presents computational as well as computation-time challenges. For example, a 64-channel EEG dataset with 200 embeddings would produce a covariance matrix of size 12,800 × 12,800. Computing the inverse and eigendecomposition of such a large dense matrix can lead to inaccuracies as well as being prohibitively slow. Furthermore, for typical EEG applications, the rank of the data is r<M, resulting from preprocessing strategies such as removing non-physiological independent components and average referencing. The first stage of source separation alleviates both of these concerns by using an optimized dimensionality reduction prior to delay-embedding.

### Selecting data for matrices S and R

“Guided” source separation methods like generalized eigendecomposition are based on a direct comparison between two researcher-selected features of the data (Parra and Sajda, 2003). Therefore, the validity and interpretability of the decomposition rests on an appropriate selection of subsets of the data from which the two covariance matrices are formed. These two covariance matrices should be similar in as many respects as possible, differing only in the characteristics that one wishes to separate. For this reason, the signal-to-noise characteristics should be similar, and the data subsets should contain a similar number of time points and trials.

For task-related designs, it is likely that **S** and **R** would come from the experimental (**S**) and control (**R**) conditions, or perhaps from all conditions combined (S) and the pre-trial baseline time period (**R**). For example, during a working memory task, the data subsets could come from the delay (memory maintenance) period and the inter-trial interval. See (de Cheveigné and Parra, 2014; Cohen and Gulbinaite, 2017) for additional discussions about data selection considerations.

### Simulated EEG data

The general procedures for simulating the EEG data will first be described, and then the specific features of the first and second simulations will be detailed (see also Figure S1 for images of key parts of the simulation process). A leadfield (anatomical forward model) was computed using OpenMEEG (Gramfort et al., 2010) as implemented in the Brainstorm toolbox (Tadel et al., 2011) in Matlab. The leadfield contains 2,004 dipoles placed in gray matter extracted from the standard template MNI brain. Each brain location was initially modeled using three dipoles for three cardinal orthogonal orientations, and these were collapsed to produce a normal vector (with respect to the cortical sheet) at each location.

Correlated random data were simulated in 2,004 dipoles as follows. First, a dipole-by-dipole matrix of positive values between 0 and 1 were computed, and this matrix was multiplied by its transpose to obtain a symmetric positive-definite matrix. The matrix values were then scaled so that the largest values were .8, except for the diagonal, which was set to 1. This matrix became the correlation matrix for all dipole time series. The next step was to simulate a 1/f power spectrum. This was achieved by scaling random complex numbers by a negative exponential to create the 1/f shape. A copy of these scaled complex numbers was then flipped to create a symmetric Fourier spectrum. The inverse Fourier transform of these values produces 1/f noise, and the real part of that result was taken. This was done for all dipoles. Next, the previously constructed correlation matrix was imposed on these data using the following formula.

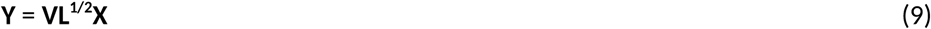

**Y** is the new correlated time series, **X** is the channels × time random number matrix, **V** are the eigenvalues of the correlation coefficient matrix, **L** contains the eigenvalues of the correlation matrix, and ^1/2^ indicates the matrix square root. Finally, the data were multiplied by 500, which is an arbitrary scaling factor selected to produce larger noise amplitude than signal.

After the dipole time series were generated, they were projected onto 64 virtual EEG electrodes arranged according to the 10-20 system and epoched into 200 trials of two seconds per trial using a sampling rate of 1024 Hz.

Additional details for the first simulation: One dipole was selected to contain the “signal” and a second dipole was selected to contain the “distractor.” The signal was a two-cycle sine wave at 5 Hz and the distractor was a three-cycle sine wave at 12 Hz. For each of the first 100 trials, the signal was summed on top of the random time series data in a randomly selected time window. The distractor was placed in all 200 trials. For convenience, the first 100 trials are referred to as “condition A” and the next 100 trials are referred to as “condition B.” The purpose of the distractor was to test whether the source separation procedure would ignore this feature of the data (it should in theory, because that feature is present in both conditions and thus does not contribute to maximizing the power ratio between *w*^*T*^***S**w* and *w*^*T*^***R**w*). Note that the signal and distractor were added to the dipole time series before projecting to the electrodes; they were not added to the electrode data.

Additional details for the second simulation: This simulation was the same as the first, except that the 12-Hz signal was added only to the first 100 trials, thus making two important features of the data that should distinguish conditions A and B. The purpose of this was to determine whether the source separation could identify and isolate both features, whether one feature would be missed, or whether both features would be mixed in the same component.

### Empirical EEG datasets

Empirical data provide important proof-of-principle applications in the context of realistic sources of signal and noise. Data used here were re-analyzed from Cohen (2015); the task is summarized here, and readers are referred to the original publication for additional details. Thirty human volunteers participated in the EEG experiment (informed consent was obtained and the study was approved by the ethics committee at the University of Amsterdam, psychology department). The task was a modified Flankers task (Appelbaum et al., 2011), in which subjects reported via button press the identity of a centrally presented letter that was flanked on both sides by other letters, which could be congruent (e.g., “T T T T T”), partially incongruent (e.g., “T T I I I”), or fully incongruent (e.g., “T T I T T”). Data were recorded from 64 electrodes placed according to the 10-20 system using BioSemi hardware (see www.biosemi.com for hardware details), sampled at 512 Hz. Additional electrodes were placed on the thumb muscles used to indicate responses; these electrodes measured the electromyogram (EMG), which was used to identify “partial errors.” Partial errors occur when subjects twitch the hand corresponding to the incorrect response but then press the correct button with the other hand. Trials containing partial errors are the strongest indicators of response conflict and elicit maximal midfrontal theta power (Cohen and van Gaal, 2014). Five trial types were separated in this task: congruent trials (the baseline condition used to create the **R** matrix), partial incongruent, full incongruent, partial errors (the condition used to create the **S** matrix), and response errors. Across subjects, the average numbers (standard deviations) of trials for these conditions, respectively, were 375 (52), 335 (54), 135 (39), 235 (99), and 84 (44). Prior to analyses, data were high-pass filtered at 2 Hz, epoched around stimulus onset, and manually inspected for excessive noise or artifacts. Data were further cleaned by removing independent components that captured oculomotor or other artifacts using the eeglab toolbox (Delorme and Makeig, 2004)(mean/std: 2.52/1.28 components per subject removed). Three datasets were excluded due to excessive noise in the data, thus the results shown here are taken from 27 individuals. Cleaner data facilitates a better decomposition, which motivated the removal of independent components and high-pass filtering.

Data for the **S** and **R** covariance matrices were taken from 0 to 600 ms post-stimulus onset. However, stimulus onsets evoke a transient phase-locked response in the EEG that can interfere with analyses of oscillatory dynamics that might co-occur with the transients, potentially leading to artifacts or misinterpretations (Yeung et al., 2007). The approach taken here to avoid potential interference from stimulus transients was to remove the phase-locked part of the signal prior to analyses (Cohen and Donner, 2013). This was accomplished by subtracting the time-domain trial average (the event-related potential) from the single-trial data, separately per condition, per channel, and per subject. The interpretation of this subtraction is that the residual—the non-phase-locked part of the signal used in analyses—can only reflect amplitude modulations of ongoing dynamics, as opposed to phase-reset transients.

Considerable previous research suggests that action monitoring tasks like this one should be associated with non-phase-locked increases in theta band (~6 Hz) activity, centered at midfrontal electrodes (around FCz or Cz), during high-conflict and error trials compared to low-conflict trials. Thus, although the “ground truth” in empirical data is not known, the expectation is that midfrontal theta should emerge as the feature of the data that most strongly separates response conflict from control conditions.

### Statistical evaluations

It is important to be aware that any statistical test between the source time series from conditions providing the **S** and **R** matrices is biased. In effect, the spatiotemporal filter is specifically constructed to maximize any possible differences between the two conditions; even with pure noise the filter will produce some result. Thus, there is a danger of overfitting, which could lead to circular inference if the results are not appropriately interpreted.

There are several approaches to address this situation. One is to apply the spatiotemporal filter to different data from those with which the filter was created. This is illustrated in the empirical data here by constructing the filter based on conditions A and D, and applying the filter to data from conditions A, B, C, D, and E. In this case, the direct comparison of D>A could be biased by overfitting, but other comparisons are not biased. Cross-validation could also be applied, in which the spatial filter is based on *N*-n trials and then applied to the remaining *n* trials. This procedure could be used to compute confidence intervals. Finally, one could use permutation testing, whereby trials within the two conditions are randomly shuffled, and many random permutations would produce a null-hypothesis distribution against which to compare the observed differences.

### Data and code availability

Matlab code to generate simulated data and apply the method is available at mikexcohen.com/data. Readers are encouraged to explore and extend the code to determine applicability of the method to their own data, as well as to test extreme and potential failure conditions.

## Results

### Simulated data

Data were created by projecting simulated dipole time series to virtual EEG electrodes, and performing all analyses on the electrode data. The first simulation involved two dipoles containing signals (brief sine waves summed on top of noise), but with only one dipole containing a “task-relevant” signal, meaning the two-cycle 5 Hz oscillation was present only in the first 100 trials (“condition A” in Figure 2). The second dipole had a three-cycle 12 Hz oscillation in both groups of trials. This second dipole acted as an irrelevant “distractor” to test the specificity of the source separation.

**Figure 2.**
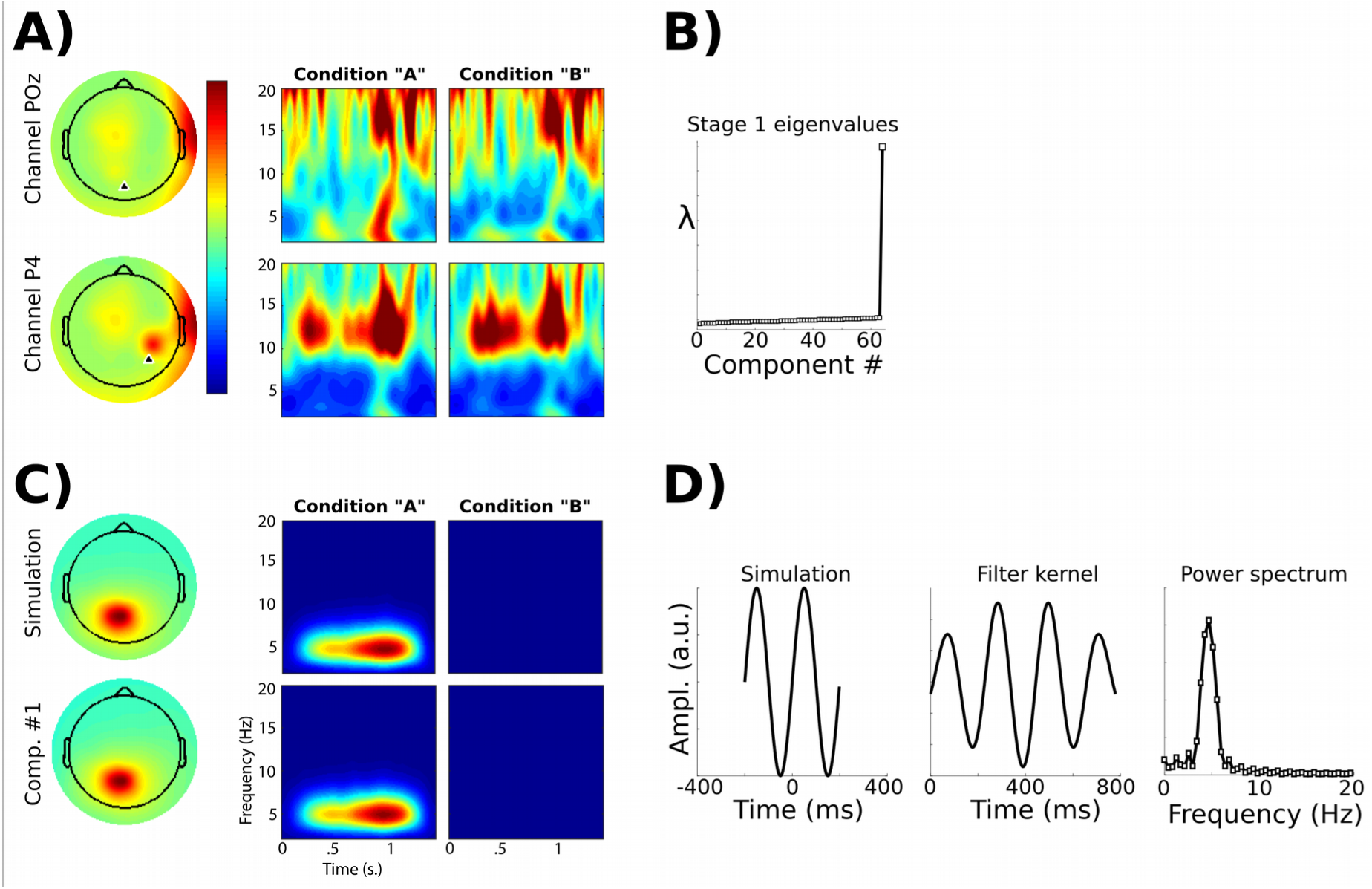
Results of the first simulation. A) Analyses of electrode-level data. The upper topographical plot depicts the spatial distribution of 5 Hz power and the lower topographical plot depicts that of 12 Hz power. The time-frequency power plots show dynamics from two electrodes (see black triangles in topographical maps) based on their proximity to the maximal projection of the dipoles selected for the simulated signals. Note that the simulated 5 Hz power is not observed due to large-amplitude noise. The two columns of time-frequency power plots correspond to condition “A” (with the 5 Hz signal) and condition “B” (without the signal). B) The stage-1 spatial source separation based on covariance matrices from conditions “A’ and “B” yielded one major component, as evidenced by a single large eigenvalue (there were 64 simulated EEG electrodes, thus producing 64 eigenvectors). C) The topographical projection of the simulated dipole and the time-frequency power plot of its time series (top row; this is “ground truth” data) and the topography and time-frequency power of the largest component. Note the similarities between the component and the ground-truth data, and their collective dissimilarity with the electrode-level results in panel A. D) Results of the stage-2 temporal source separation. The left plot shows the simulated signal. The middle plot shows the temporal filter kernel (cf. Figure 1i), and the right plot shows its power spectrum.

Electrode-level analyses were unable to identify the 5-Hz signal, because its amplitude was comparable to the noise level. The 12-Hz “distractor” was visible, because its source amplitude was higher than that of the noise. The spatial source separation (steps a-e in Figure 1) on the covariance matrices comparing conditions A and B recovered the spatial topography as well as the time-frequency characteristics of the signal. The second source separation stage recovered an empirical filter kernel that had a similar shape and spectral profile as the original simulated data (Figure 2D).

In the second simulation, the two dipoles contained “task-relevant” signals, with one having an oscillation at 5 Hz and the other at 12 Hz. The purpose of this simulation was to test how two components would be identified by the spatiotemporal decomposition, considering that both features are task-relevant.

Results showed that the two spectral-spatial features were isolated into different components. This can be seen by two relatively large eigenvalues from the first stage of source separation (Figure 3b). The associated eigenvectors isolated spatial components that were consistent with the topographical projections of the two dipoles (Figure 3c,e). The second stage of source separation was performed on two separate Hankel matrices: one created from the time series of the largest component, and one created from the time series of the second-largest component. The resulting temporal filters accurately reconstructed the spectral characteristics of the two simulated time series (Figure 3d,f). The electrode-level analyses partially revealed the simulated data, but were also considerably noisier. Without *a priori* knowledge of the simulated data, it would be difficult to know which time-frequency-electrode features reflect “true” signals. Overall, results from this simulation confirm that it is possible to separate multiple narrowband spatial-temporal components in multichannel data without applying any narrowband temporal filters.

**Figure 3.**
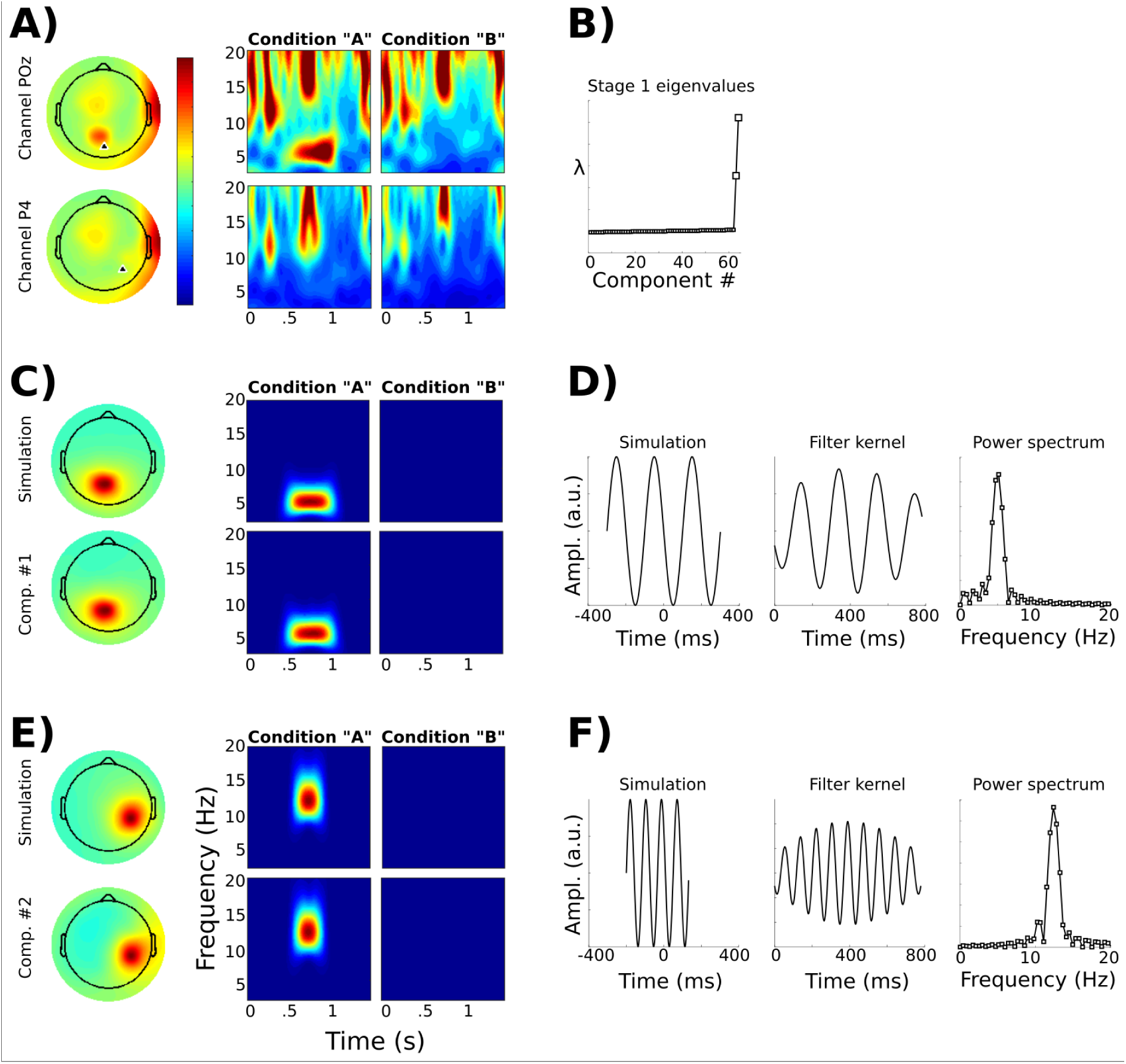
Results of the second simulation. This simulation was similar to the first with the addition of a second task-related signal in a second dipole at 12 Hz. This figure is organized similarly to Figure 2. A) Electrode-level data. B) Note that the stage-1 source separation revealed two spatial components with relatively large eigenvalues. The largest component isolated the 5 Hz signal while the second component isolated the 12 Hz signal (the 5 Hz component was larger because the signal time series was longer). Note that despite the two signals overlapping in time and in topography, they are fully isolated into two distinct components because their trial-to-trial temporal onsets were non-phase-locked, thus allowing sufficient spatial-temporal separation. No narrowband filters were applied in either of the two source separation stages (narrowband filters were applied only to obtain the time-frequency power plots).

A third simulation (using only the second stage on single-channel time series data) was conducted to illustrate how the empirical filter kernel identifies the most prominent features of the data that distinguish it from the reference time series, which may not capture all subtle features of the waveform shape. A square wave with a linear trend was added to random white noise (see Figure 4a for the simulated signal and an example single trial of the signal plus noise). The reference time series was noise. The filter kernel had a sinusoidal shape, which captured the most distinctive temporal feature relative to the reference (note that this is not necessarily the same as the most visually salient feature of the simulated waveform). This empirical filter kernel was then applied to the time series data in Figure 4a (this would be the “measured” data), revealing the rank-1 approximation of the signal that best separates the signal from the reference data. Although the reconstruction does not capture the high-frequency waveform features such as sharp edges, it represents the temporal features that best distinguish the **S** from **R** time windows.

**Figure 4.**
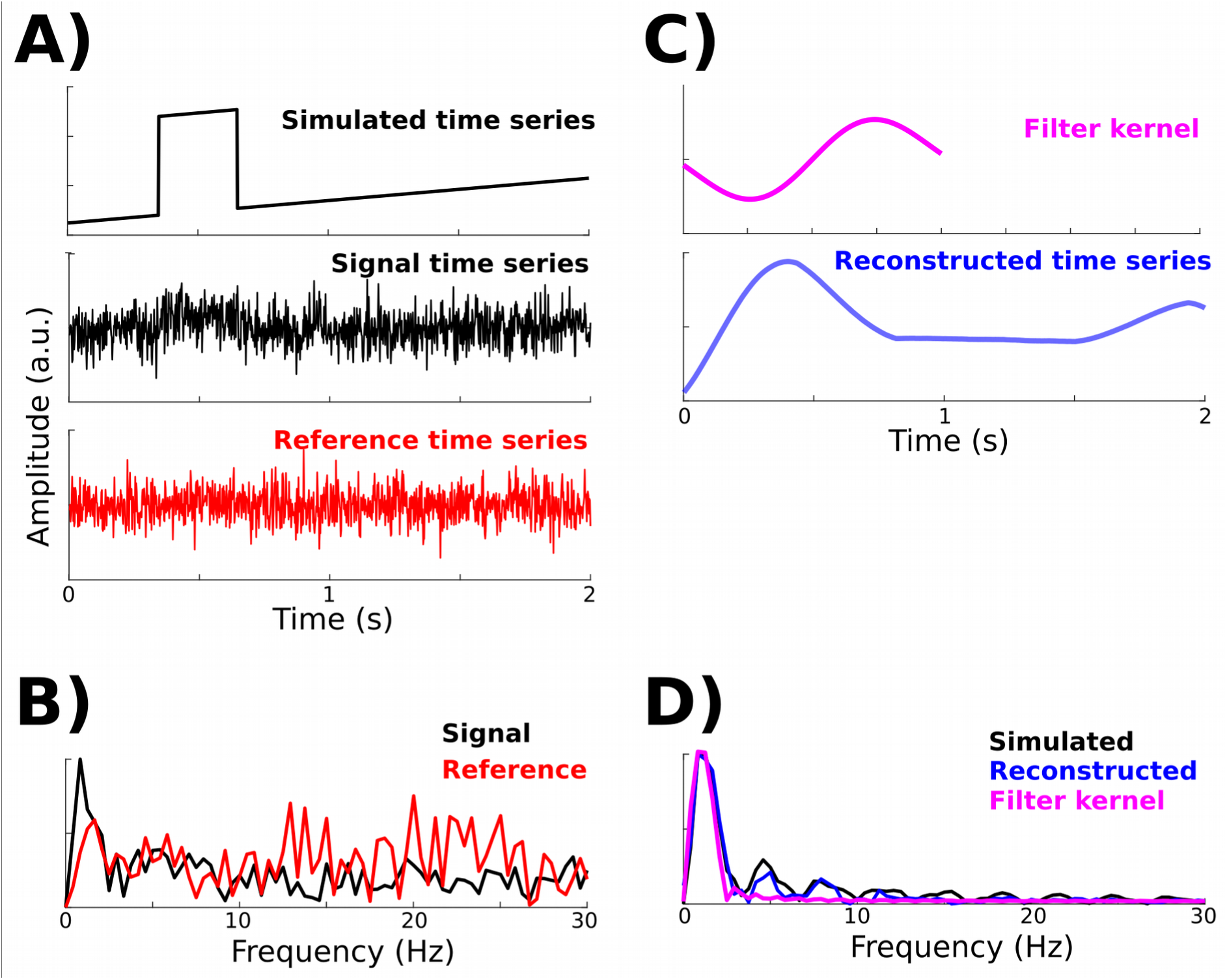
Simulation of non-stationary time series. (A) The simulated (ground-truth) data and examples of two single trials used to construct the **S** matrix (containing signal and noise) and the **R** matrix (containing only noise) (the top plot has a different y-axis scaling for visibility). (B) The power spectra from these two example trials. (C) The empirical temporal filter that maximally separated **S** from **R** was concentrated in the low-frequency range (although the filter was not based on frequency-domain or filtered data). The reconstructed single-trial signal is smooth relative to the simulated time series, but is a better approximation than the “measured” data in the middle row of panel A. (D) The power spectra of the simulated signal, empirical kernel, and reconstructed signal.

### Empirical data

The procedure outlined in Figure 1 was applied to empirical EEG data. The dataset had five experiment conditions related to response conflict, corresponding to a baseline (no response conflict), three levels of response conflict during correct trials, and response errors. Both source separation stages were based on comparing the condition with the strongest response conflict (correct trials containing partial errors) with the baseline condition (congruent trials). After the spatiotemporal filters were constructed based on these conditions, they were applied to all five conditions.

Figure 5 shows the group-average topographical projection of the spatial filter, the spatiotemporal filter kernel, its power spectrum, and the power envelope computed as the squared magnitude of the Hilbert transform applied to the spatiotemporal component. Figure 5 shows the topographical projections, the time-domain filter kernel projection, and its power spectrum, for each individual subject.

**Figure 5.**
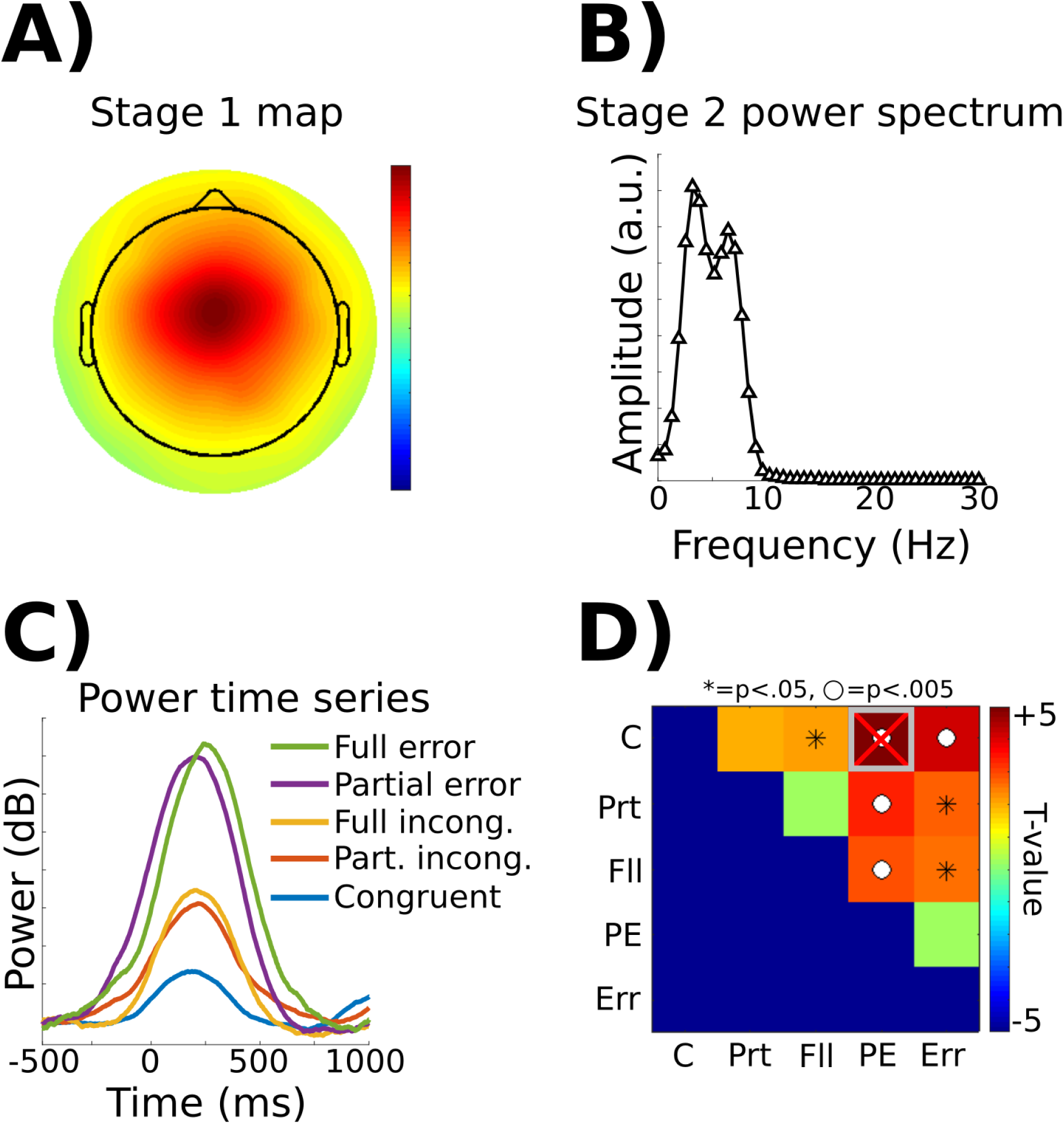
Group-level results of the spatiotemporal filter on empirical EEG data (N=27 humans). The **S** and **R** matrices were generated, respectively, from conditions with high vs. low response conflict (“Partial error” and “Congruent”). (A) The stage-1 maps indicated a midfrontal-focused component. (B) The power spectrum of the temporal filter had peaks at 3.2 and 6.4 Hz. This apparent double peak resulted from averaging individual narrow peaks, as can be seen in Figure 6. (C) The baseline normalized power time series (extracted from the squared magnitude of the Hilbert transform of the stage-2 component time series) showed a peak at around 250 ms. (D) Average power from 0-600 ms was used for an all-to-all t-test matrix. The Bonferroni-corrected threshold of p<.05/10 as well as the uncorrected p<.05 threshold results are indicated. The comparison between partial errors and congruent trials is biased because these are the conditions used to define the spatiotemporal filter; this result should be interpreted with caution. (C=congruent, Prt=partial conflict, Fll=full conflict, PE=partial error, Err=full error).

Several aspects of these results are worth remarking. First, because the phase-locked (ERP) component of the signal was removed prior to analyses, these results reflect only non-phase-locked dynamics and are not influenced by phase-locked or evoked transients. Second, the optimal spatiotemporal filter was narrowband, despite the complete absence of any narrowband filters applied to the data. This demonstrates that the narrowband activity was endogenously present in the data, and not imposed by narrowband filtering a non-oscillatory evoked response, as has been suggested could occur (Yeung et al., 2007). Third, although the difference between partial error and congruent trials can be expected based on overfitting noise (both source-separation stages were based on separating these two conditions), the differences for other conditions are not trivial, as those data were not considered when constructing the filters.

Finally, it is interesting to inspect the individual variability in the topography and frequency of the spatiotemporal feature that best distinguished response conflict from the congruent condition (Figure 6). The origin of this variability is not further investigated here, but it is possible that these differences are related to meaningful variability in genetics, age, or brain structure (Klimesch, 1999: Haegens et al., 2014: Cecere et al., 2015). Two subjects had stage-1 topographical projections suggestive of artifacts (7th in the first column and 2nd in the second column of Figure 6). Closer inspection of the data, however, did not reveal excessively noisy or corrupted data, and there was no clear justification for removing these datasets from the analyses.

**Figure 6.**
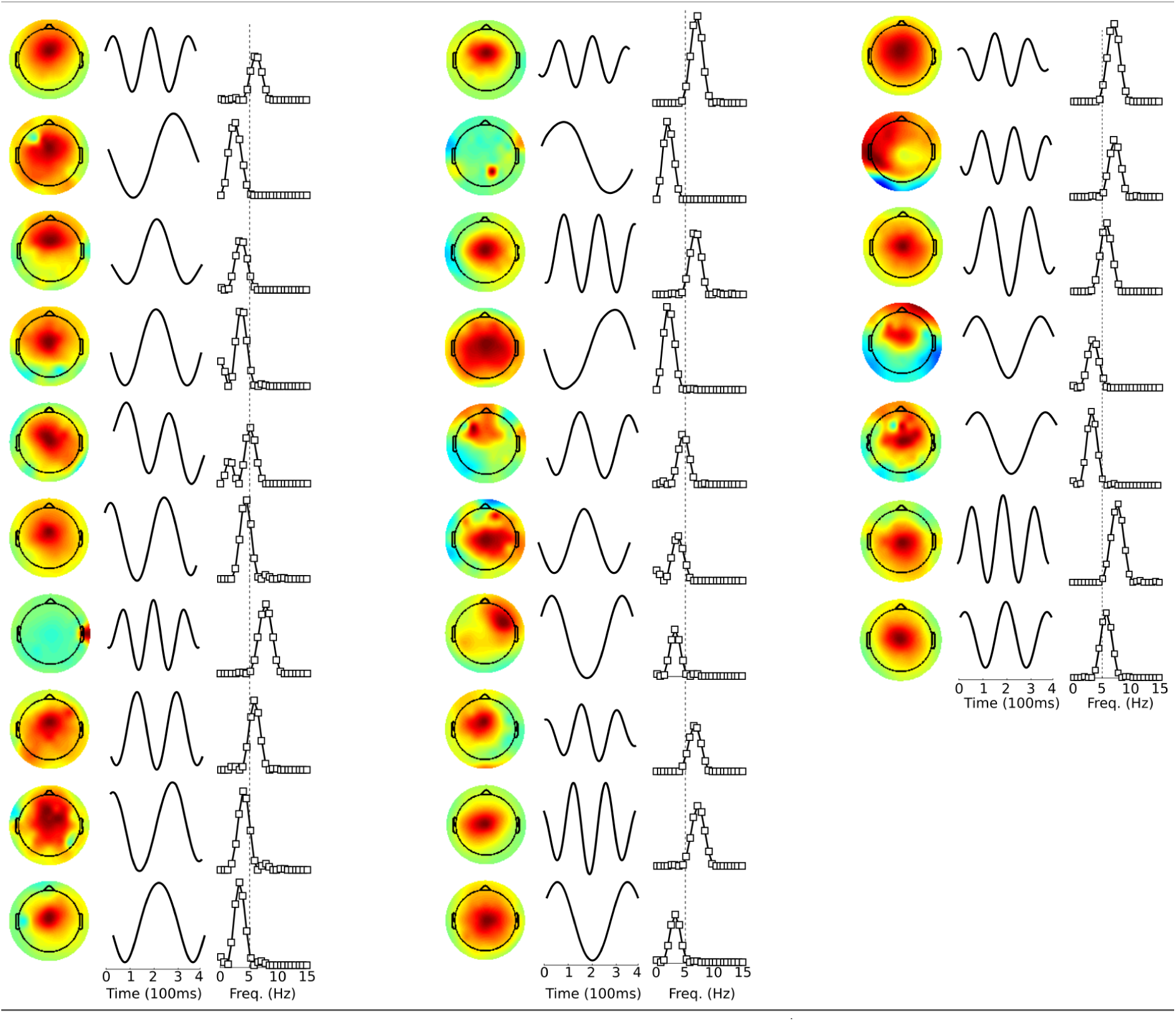
Individual data for all subjects from the experiment (the ordering is based on data acquisition date and is therefore arbitrary with respect to the results). These topographical maps were averaged together in Figure 5a, and the power spectra were averaged together in Figure 5b. The vertical dashed line indicates 5 Hz for reference. Note that each individual subject had a narrow peak, but variability in the peak frequency led to the apparent double-peak in Figure 5b. The time courses show the stage-2 source separation filters. No narrowband filters were applied at any stage; these signal characteristics were empirically identified by the decomposition as being the most relevant features for distinguishing partial error from congruent trials.

## Discussion

Population-level neural activity is often rhythmic, and these rhythmic patterns are increasingly being linked to healthy and to dysfunctional cognitive and perceptual processes. Important insights into the relationship between rhythmic neural activity and brain function will come from understanding the neurophysiological principles that produce these rhythms, and how those principles are related to the neural computations that implement cognitive operations. This endeavor is complicated by several limitations, such as large noise relative to signal (which is generally worse for non-invasive measurements) and each electrode measuring activity simultaneously from multiple sources of signal and noise. Multichannel recordings can help ameliorate these limitations, because the different sources of activity project instantaneously and linearly onto different electrodes. This fact helps source-separation techniques recover the underlying sources, assuming the statistical features of the sources conform to the assumptions made by the source separation method applied.

Most existing source-separation methods focus exclusively on optimizing spatial (electrode) weights, while using traditional (e.g., Fourier-based) signal processing tools for subsequent temporal analyses. This paper showed that the same source separation techniques can be applied to univariate time series data as well, with the goal of empirically identifying temporal patterns that discriminate between two conditions or two time windows. One advantage of this method is that it eliminates the need to impose temporal filters with specified temporal structures (such as sine waves), which may be unrelated to the temporal process that generates the measured activity. This is not to say that traditional temporal signal processing methods are inappropriate; instead, it is important to have many tools in a scientist’s toolkit.

### Advantages and limitations

Source separation methods in general have several advantages. They increase the signal-to-noise characteristics, they help identify patterns in the data that might be difficult to obtain from single-electrode analyses, they reduce the dimensionality of the data in a “guided” way (in contrast to completely blind decompositions), and they reduce the need for potentially suboptimal electrode selection (Makeig et al., 2004; Blankertz et al., 2008; Cunningham and Yu, 2014; Cohen, 2016). The extension to temporal source separation illustrated here provides additional benefits, including blind discovery of prominent temporal characteristics that can be used for empirically derived filter kernels, and further separating signal from noise in time series data.

The second stage of source separation does not require multichannel data; it could be performed on single-channel time series. The advantage of the first stage is to facilitate isolation of a single spatial component. This can be particularly important for datasets in which multiple sources contribute to the data recorded at each electrode. In these cases, the first source separation phase will help spatially isolate features of the data, which will facilitate the temporal separation at the second stage. It would also be possible to apply temporal filters to the data prior to the stage-1 source separation (Cohen, 2016). However, such temporal filters should be fairly wide, otherwise the stage-2 temporal filter may simply reflect the sinusoidality imposed by the narrowband filter.

### Empirical temporal filter kernel versus waveform shape

Perhaps the main limitation of the method presented here is related to the subtle but important distinction between the waveform shape that reflects the biophysics of the neural circuit that produces measurable electrical fields, vs. the empirically derived temporal filter kernel obtained from the method described here. The waveform shape of brain oscillations has become increasingly a topic of conversation in the neuroscience literature (Jensen et al., 2010; Jones, 2016; Cole and Voytek, 2017). Identifying waveform shape is important because it provides an anchor-point for linking EEG results to underlying neurophysiology. Many empirically measured waveforms appear sinusoidal, but this may result from using sinusoidal filter banks to identify those waveforms; thus, the sinusoidal filters will identify only the sinusoidal features of the “true” waveform. There are a few striking cases of neural oscillations having non-sinusoidal shapes (Cole and Voytek, 2017), but these tend to be unusually strong signals measured invasively, such as rat hippocampal theta.

The source separation method presented here produces the low-rank approximation of the time series that optimally achieves the specified constraints (e.g., difference between conditions or time windows). This is not the same as the waveform shape itself. For example, features such as a sharp edge or a small ripple may be visually salient and physiologically meaningful, but if they contain little discriminative information, those features will be ignored by source-separation or machine-learning algorithms. Indeed, eigendecomposition and related methods have the goal of optimizing subspace basis vectors based on patterns in covariance matrices; they are not constrained by potential neurophysiological plausibility. Although this hinders a simple physiological interpretation, it is also an advantage: One need not specify a large number of unknown parameters and constraints for the method to be valid and appropriately used. Therefore, if the empirical filter kernel is narrowband, it indicates that a narrowband feature best discriminates two conditions, although it does not mean that additional features of the data are irrelevant.

### Automatic or manual component selection?

In theory, algorithmic component selection would be optimal because it eliminates the potential for researcher bias or subjectivity. The easiest selection criteria would be to take the eigenvector with the largest corresponding eigenvalue. Additional selection criteria can be incorporated based on *a priori* expectations of the results, such as maximal topographical projection onto some electrodes, or maximal spectral power within some frequency range.

However, a simple selection algorithm may not always select the most appropriate component. Therefore, some expert user selection may be necessary. This is analogous to user-guided selection of components during independent components analysis. Indeed, algorithm-based selection methods of independent components tend to be suboptimal relative to expert human selection (Chaumon et al., 2015). Human-supervised component selection should not be avoided or shunned. As long as the component is selected in a way that is orthogonal to the main analysis, the risk of introducing systematic biases towards any particular statistical result can be minimized.

### Comparison to other source separation methods

There are many source separation methods that range in assumptions and implementation details. The brain-computer-interface community has developed many strategies for dimensionality reduction and source separation as it relates to classification of states based on multichannel EEG signals (Fouad et al., 2014). Generalized eigendecomposition is used in many of these approaches (where it is sometimes called common spatial pattern analysis), because it tends to be a fast, robust, and efficient method.

The primary novelty of the present paper is to demonstrate that the same source separation principle can be applied to time series data by first expanding the dimensionality of a time series using delay-embedding. The use of delay-embedded matrices in neuroscience is already established. For example, Brunton et al. (2016) used delay-embedded matrices to estimate data components (spatiotemporal coefficients) that link data at each time point to data at the previous time point. Lainscsek and Sejnowski (Lainscsek and Sejnowski, 2015; Lainscsek et al., 2015) used delay-embedded matrices to model neural dynamics as delay differential equations to estimate frequency-specific responses and couplings between electrodes. Delay-embedding matrices are a powerful method for uncovering dynamics in time series data, and continued methodological development and applications will improve the quality of neuroscience data analysis.

An advantage of using generalized eigendecomposition for the temporal source separation is that, like with the spatial source separation, it is fast and robust, and does not require any parameters other than those used to select the data for the **S** and **R** matrices. Applying the source separation in two steps (first spatial, then temporal) is a major advantage. As mentioned in the Methods section, trying to combine these into a single analysis step is theoretically sensible, but practically difficult due to instabilities of decompositions on very large spatiotemporal matrices.

### Implications for midfrontal theta and response conflict

Midfrontal theta is a robust neural signature of action monitoring including response conflict, error detection, and learning from negative feedback (Cavanagh and Frank, 2014; Cohen, 2014b). Because the primary purpose of this paper is methodological, there are limited novel insights into the neural mechanisms of response conflict processing and performance monitoring. That said, this study provided an independent demonstration using a novel analysis technique that conflict-related midfrontal theta reflects an amplitude modulation of ongoing theta oscillations, as opposed to a phasic non-oscillatory evoked potential (for further discussion of this point, see Yeung et al., 2004, 2007; Trujillo and Allen, 2007; Cohen and Donner, 2013; Munneke et al., 2015).

## Conclusion

Multivariate source separation methods are becoming increasingly important in neuroscience, as the number of simultaneously recorded electrodes is steadily increasing (Stevenson and Kording, 2011) and as it is increasingly becoming clear that information can be embedded within spatial-temporal patterns of data that may be difficult to ascertain from traditional (e.g., single-channel ERP) analyses. Although source-separation methods are typically applied only in the spatial dimension, they can also be applied directly to the time series data to create empirical temporal filters that maximize researcher-defined criteria without the necessity to apply sine wave-based filters that may distort non-stationary time series. This was illustrated here by applying generalized eigendecomposition to delay-embedded matrices. In the empirical EEG application, it was demonstrated that response conflict and errors elicit non-phase-locked narrow-band activity recorded over midfrontal regions. The consistency of this finding with previous studies proves a proof-of-principle demonstration of the method, as well as confirming the role of theta oscillations in cognitive control processes without the necessity to apply narrowband sinusoidal filters.

---

**Figure S1.**
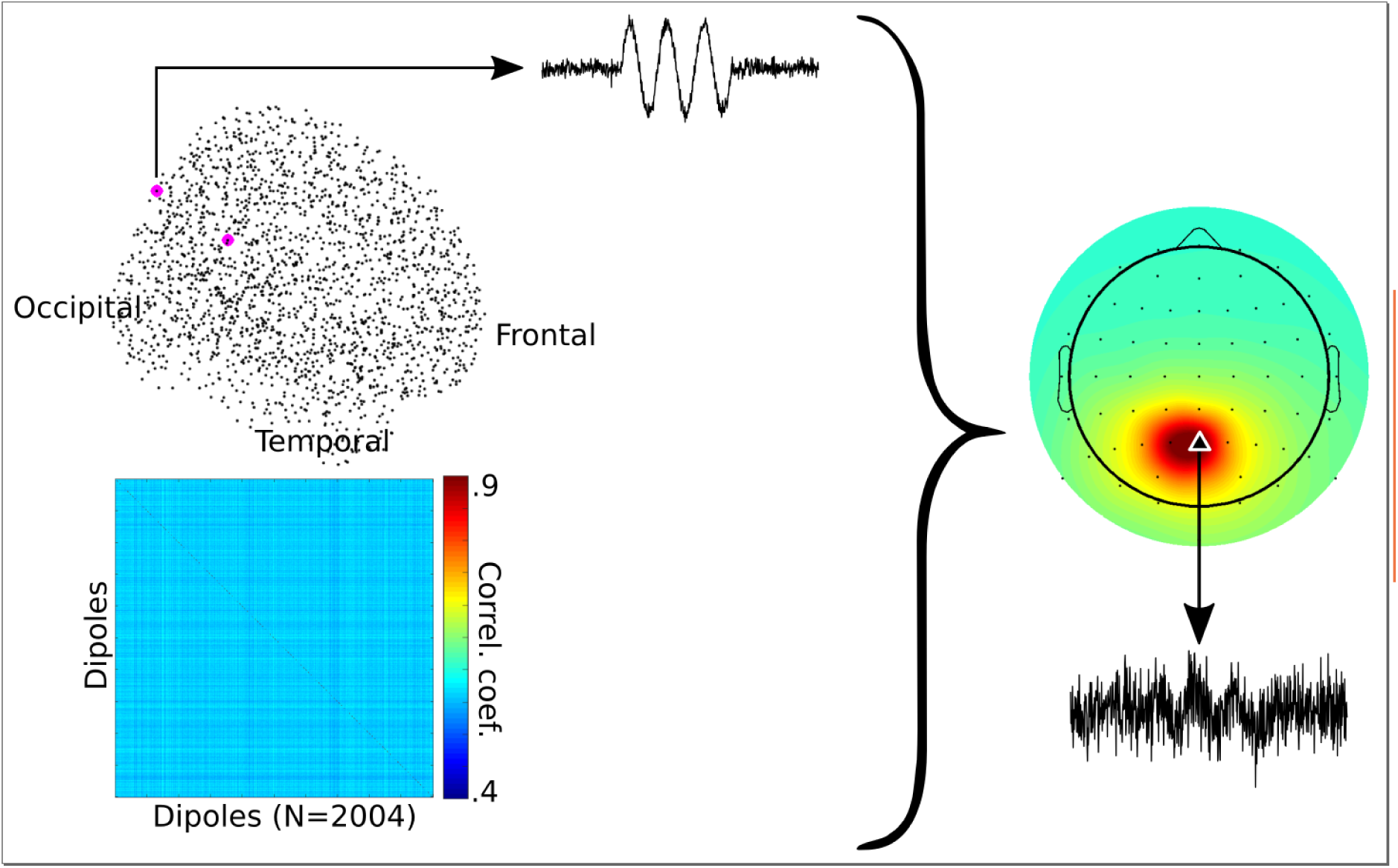
Overview of key elements of data simulation. 2,004 dipoles were placed in the cortex (black dots) with an orientation normal to the cortical surface. Noise data were generated by imposing a correlation structure (see correlation matrix) on random numbers that had a 1/f power spectrum. Two dipoles (magenta) were selected to contain brief sine waves that were summed on top of the noise. Finally, the time series from all dipoles were projected onto the scalp and summed. Note the difference in signal amplitude from the dipole to the EEG electrode with maximum dipole projection; this difference is due to source-level mixing with activity from other dipoles.

